# Seasonal H1N1 influenza A virus preferentially infects a pre-existing ciliogenesis-primed secretory cell state

**DOI:** 10.64898/2026.11.04.764425

**Authors:** Josephine von Kempis, Peter Leary, Lennart Opitz, Silke Stertz

## Abstract

Influenza A viruses (IAV) infect secretory and ciliated cells of the human airway epithelium, but whether all cells within one cell type are equally susceptible to infection remains unresolved. Here, single-cell RNA sequencing analysis of primary human bronchial epithelial cultures from three donors identified preferential infection of a transcriptionally distinct subpopulation of secretory cells by seasonal H1N1 IAV A/Hawaii/70/2019. Genes upregulated in this subpopulation showed enrichment for cilium-related gene ontology terms. In an *ex post facto* approach, this enrichment persisted after excluding genes differentially expressed between infected and mock-treated samples, including those identified in secretory cell-specific comparisons. Single-cell analysis further revealed no significant increase in ciliogenesis scores in infected secretory cells compared to mock infected cells at 6 hours post-infection, arguing against infection-induced activation of the canonical ciliogenesis programme during the initial viral replication cycle. Collectively, these results support preferential infection of ciliogenesis-primed secretory cells and refine IAV tropism beyond cell type to encompass epithelial differentiation state.

## INTRODUCTION

Despite the availability of vaccines and antiviral therapies, influenza remains a major global public health challenge (*1*). Influenza is an acute respiratory disease commonly known as the flu and caused primarily by influenza A viruses (IAVs). According to the World Health Organization (WHO), these circulating viruses account for an estimated one billion infections, three to five million cases of severe illnesses, and 290,000 to 650,000 deaths annually worldwide. Beyond the substantial health impact, influenza provokes a significant economic burden, with annual costs estimated at six to fourteen billion euros in the European Union alone (*2*).

Due to its ability to spread via airborne droplets and aerosols, IAV enters its host through the respiratory tract, making the upper airway the initial site of viral attachment and infection (reviewed in (*3*)). This respiratory tissue comprises a ciliated pseudostratified columnar epithelium that serves as the first barrier to potential pathogens and toxins, while also being the initial target of respiratory viruses. The airway epithelium exerts its barrier function, for example, through the secretion of mucus and mucociliary clearance. In this highly structured self-retaining microenvironment, the cell layer can differentiate through its stem cell population, known as basal cells, to all other cell subtypes present in the epithelium (reviewed in (*4*)). This includes different minor or transient cell populations like ionocytes (*5*, *6*), required for ion transport across the cell layer; tuft/brush cells (*7*, *8*), specialized chemosensory cells; and neuroendocrine cells (*9*, *10*), which bridge the nervous and the endocrine systems. Next to these minor cell populations, the predominant differentiation path of basal cells involves activation of the NOTCH signalling pathway, which induces the formation of club cells (reviewed in (*11*)). Club cells are cells producing and secreting protective uteroglobin (SCGB1A1) proteins (*12*, *13*). Besides their secretory function, club cells exhibit progenitor cell properties and can differentiate into mucus-producing goblet cells (*14*) or, via a distinct pathway, into ciliated cells. The differentiation of club cells to ciliated cells is governed by the highly regulated process of ciliogenesis (reviewed in (*15*)). In brief, this transcriptional pathway is initiated by the downregulation of the transcription factors NOTCH and GEMININ (*11*, *12*, *16–18*). This downregulation consequently allows for the upregulation of GEMC1 to interact with E2F4 and E2F5, as well as the transcription factor TF-DP1 (also referred to as DP1). These interactions, in return, promote the transcriptional induction of MCIDAS, which also forms protein complexes with E2F4/5 and DP1 and ultimately supports the expression of the transcription factor TP73. TP73 is a key factor allowing the upregulation of MYB, CDC20B, and CCNO, which are involved in centriole amplification (*19–21*). The centrioles arise from DEUP1-enriched deuterosomes (*22–24*). Thus, this intermediate cell state has previously been referred to as the deuterosomal cell state (*25*). Each centriole will further mature into a basal body during the process of ciliogenesis. TP73 also enables upregulation of FOXJ1, RFX2, and RFX3. These proteins further direct the maturation and apical anchoring of the basal bodies (reviewed in (*26*)) and induce the expression of cilia structural components (such as dyneins) and therefore regulate the length of the cilia (*27*, *28*). Ultimately, the genes inducing cilia motility are upregulated and complete the ciliogenesis differentiation program (reviewed in (*29*)). Mature ciliated cells, through the coordinated beating of the motile cilia, facilitate clearance of mucus from the airway epithelium together with any trapped material or pathogen.

Recent advances in single-cell RNA sequencing have allowed more detailed analysis of the processes of cellular differentiation (*30*). These technologies have also been increasingly applied to investigate cell state dynamics in the context of viral infections (reviewed in (*31*)). One prominent example is a study by Greaney *et al.* (2023), who revealed that severe acute respiratory syndrome coronavirus 2 (SARS-CoV-2) preferentially enters ciliated precursor cells (*32*). These precursors are cells in the process of differentiating from secretory cells to mature ciliated cells. Owing to their rapid epithelial turnover and active differentiation, this precursor population may provide a favourable environment for viral replication. Greaney *et al.* (2023) further showed that the virus promoted ciliated cell differentiation and thus gave rise to a population of infected mature ciliated cells. Kim & Lee (2024) also reported that SARS-CoV-2 infects an intermediate differentiation stage between basal and ciliated cells (*33*) and thus support the finding of Greaney *et al.* (2023). In contrast to the finding of infection-induced ciliated cell maturation, Berg *et al.* (2026) reported respiratory syncytial virus (RSV) infection causing a shutdown of the ciliogenesis pathway and thus leading to ciliated cell dedifferentiation (*34*).

In von Kempis *et al.* (2026), we previously demonstrated that human influenza H1N1 virus infects both ciliated and secretory cells of the human bronchial epithelium within the first round of virus replication (*35*). This observation raised the following questions: (i) Does the seasonal influenza H1N1 virus preferentially infect secretory cells that are primed for ciliogenesis, or does the virus infect secretory cells independent of their differentiation state? (ii) Does the seasonal influenza H1N1 virus induce infection-dependent cell subtype transitions, as previously observed with SARS-CoV-2 and RSV?

In this report, using a single-cell RNA sequencing *ex post facto* approach, we reveal that during the initial infection cycle of the virus, human H1N1 seasonal 2019 IAV preferentially targets a specific bronchial secretory cell subtype characterised by elevated expression of cilium-related genes. We further provide evidence that this tropism is driven by a pre-existing secretory cell state, rather than by an infection-induced cell subtype transition to ciliated cells and thus refine IAV tropism to include the transcriptional state in addition to the cell type.

## RESULTS

### Infected secretory cells are transcriptionally distinct from bystander secretory cells

In a previous study, we acquired single-cell sequencing datasets from primary human bronchial epithelial cell cultures from three independent donors infected for 6 hours with H1N1 A/Hawaii/70/2019 virus or a mock control (*35*). From the quality-controlled and filtered dataset, data points were normalised and batch-corrected, then further dimensionally reduced and clustered based on transcriptional similarities among cells. Cell subtypes were determined by using canonical markers identified as cluster markers. We distinguished the following epithelial cell subtypes in the bronchial cultures: ciliated, secretory, intermediate, basal cells, and ionocytes. Due to the similarities of the transcriptomes from club and goblet cells, we grouped them as ‘secretory cells’. The infected cells were further defined as cells expressing transcripts from at least 4 out of the 8 viral genome segments. We concluded that the human IAV strain infected around 41% of all ciliated cells and around 20% of the secretory cell population (Supp. Fig. 1A).

To characterise the differences between infected versus bystander (non-infected) cells, we used pseudobulk expression profiles and performed differential expression analysis comparing first Hawaii/70/2019 infected ciliated cells over bystander ciliated cells (Fig. 1A). A differentially expressed gene (DEG) was defined as a gene with an absolute log2 fold change ≥ 0.5 and with a false discovery rate (FDR) ≤ 0.05. A gene ontology (GO) enrichment analysis on the hits overrepresented in the infected ciliated cell population suggests that processes related to protein translation are more prominent in the cells primed for infection (Fig. 1B-D).

**Figure 1:**
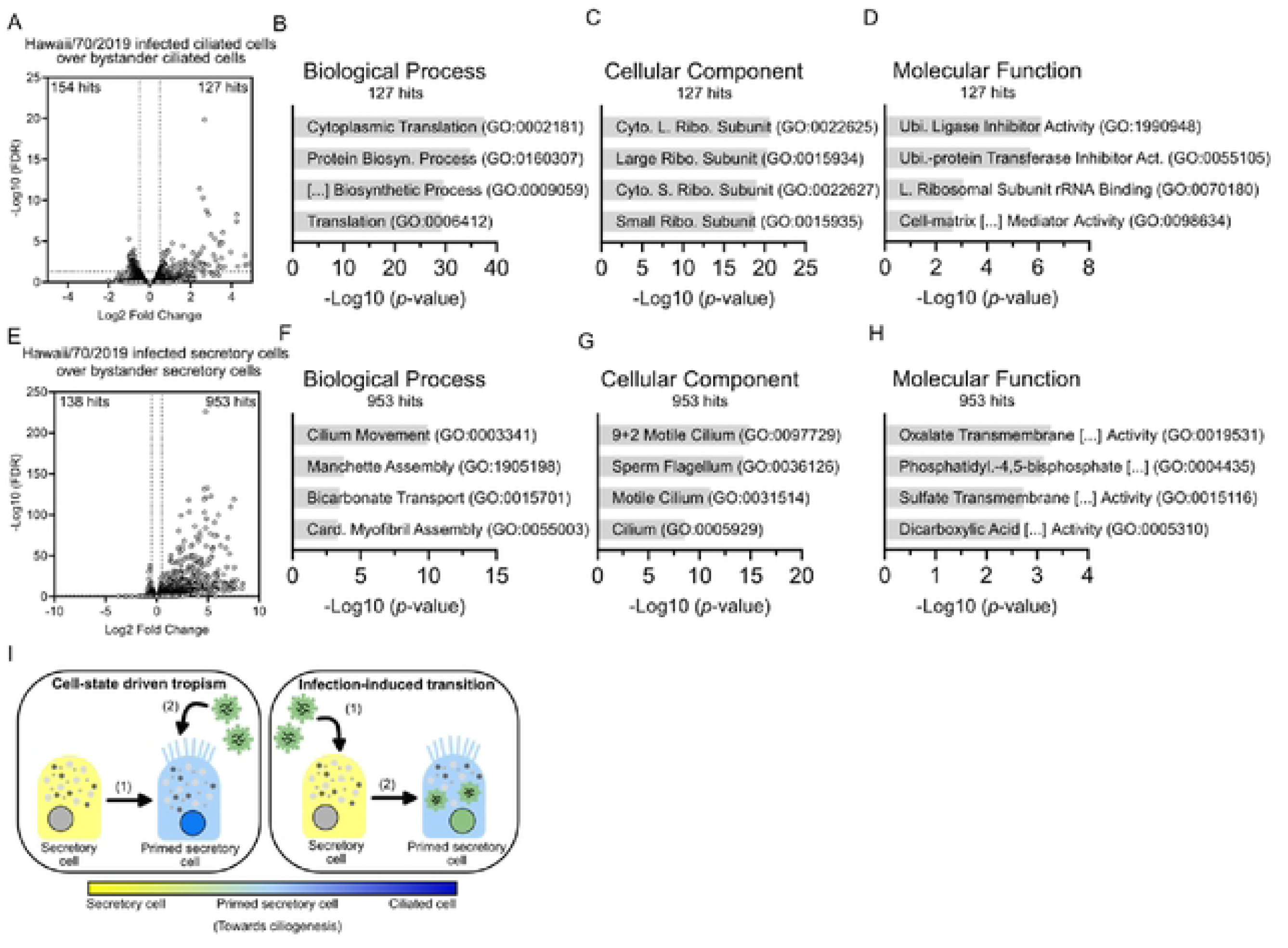
Infected secretory cells are transcriptionally distinct from bystander secretory cells. **A)** Volcano plot of scRNAseq data from BEpC from donors 1-3 infected with H1N1 Hawaii/70/2019 for 6 hours with an MOI of 1. The scRNAseq dataset (n=3) was pooled to generate pseudobulk expression profiles, and a differential expression analysis was performed of Hawaii/70/2019 infected ciliated cells (686 cells) over bystander (non-infected) ciliated cells (959 cells). **B)** - **D)** Gene ontology (GO) enrichment analysis of the hits overrepresented (Log2 (fold change) ≥ +0.5 and FDR ≥ 0.05) in the Hawaii/70/2019 infected ciliated cells over bystander ciliated cells analysis Top 4 GO terms are shown. **E)** The scRNAseq dataset was pooled (n=3) to generate pseudobulk expression profiles, and a differential expression analysis was performed of Hawaii/70/2019 infected secretory cells (828 cells) over bystander secretory cells (3538 cells). **F)** - **H)** Gene ontology (GO) enrichment analysis of the hits overrepresented (Log2 (fold change) ≥ +0.5 and FDR ≥ 0.05) in the analysis Hawaii/70/2019 infected secretory cells over bystander secretory cells. Top 4 GO terms are shown. **I)** Schematic representation of the study hypothesis.

Next, we applied the same analysis to compare infected secretory cells over bystander secretory cells (Fig. 1E). This analysis identified 953 transcripts significantly overrepresented in infected secretory cells, and thus a substantially higher number of hits compared to infected ciliated cells. In addition, the transcripts exhibited larger fold changes and lower FDR values (Fig. 1A, E), suggesting more pronounced and statistically robust transcriptional differences between infected and bystander secretory cells. The comparatively small number of differentially expressed transcripts and lower fold changes observed in infected ciliated cells suggest that infected and bystander ciliated cells have more similar transcriptomes. Given the pronounced transcriptional differences observed in infected secretory cells, we focused on this population for further characterization.

To understand the overrepresented transcripts, we performed a GO enrichment analysis on the DEGs in infected secretory cells (Fig. 1F-H). Interestingly, the most significantly enriched GO terms were strongly associated with the cilium and ciliary motility. Based on this finding, we hypothesized that secretory cells infected by the seasonal H1N1 influenza virus are transcriptionally highly distinct from bystander secretory cells and that the Hawaii/70/2019 virus preferentially infects secretory cells primed for ciliogenesis (Fig. 1I, left panel). However, if infection-induced changes were themselves the drivers of ciliogenesis in infected cells, then the data would alternatively support a model in which viral infection induces a transition of secretory cells to ciliogenesis-primed cells (Fig. 1I, right panel).

### Secretory cells are the precursors of ciliated cells

To determine whether the *in vitro* cultured bronchial airway epithelial cells differentiate comparably to *in vivo* epithelia, we conducted a trajectory inference analysis. We used the mock-treated samples to ensure that the pathway analysis is independent of any infection-induced alterations. We conducted a differentiation trajectory analysis and rooted it at the basal cell cluster 7 due to the high expression of basal canonical markers *KRT14* and *KRT5* (Fig. 2A-D). We first used the trajectory inference slingshot approach (*36*), which runs on the 30-dimensional Harmony batch-corrected embedding (Fig. 2E). The analysis modelled the first pseudotime lineage going through the basal cell populations (7 ◊ 0 ◊ 2), then through the intermediate cell population (cluster 6), and before bridging into the ciliated cell clusters (3 ◊ 5 ◊ 11), the trajectory passes by the secretory cell cluster 4. This result therefore supports the *in vivo* differentiation pathway of ciliated cells from basal via secretory cells (reviewed in (*15*)). The top hits (based on fold change) increasing from the intermediate population (cluster 6) to the ciliated cells (cluster 5) are, for example, the ciliary-associated calcium-binding coiled-coil protein 1 *CABCOCO1* and the cilia formation regulator *ERICH3* (Supp. Fig. 2A) (*37*, *38*), while two of the top decreasing transcripts are the canonical basal cell markers *KRT5* and *KRT6A* (Supp. Fig. 2B).

**Figure 2:**
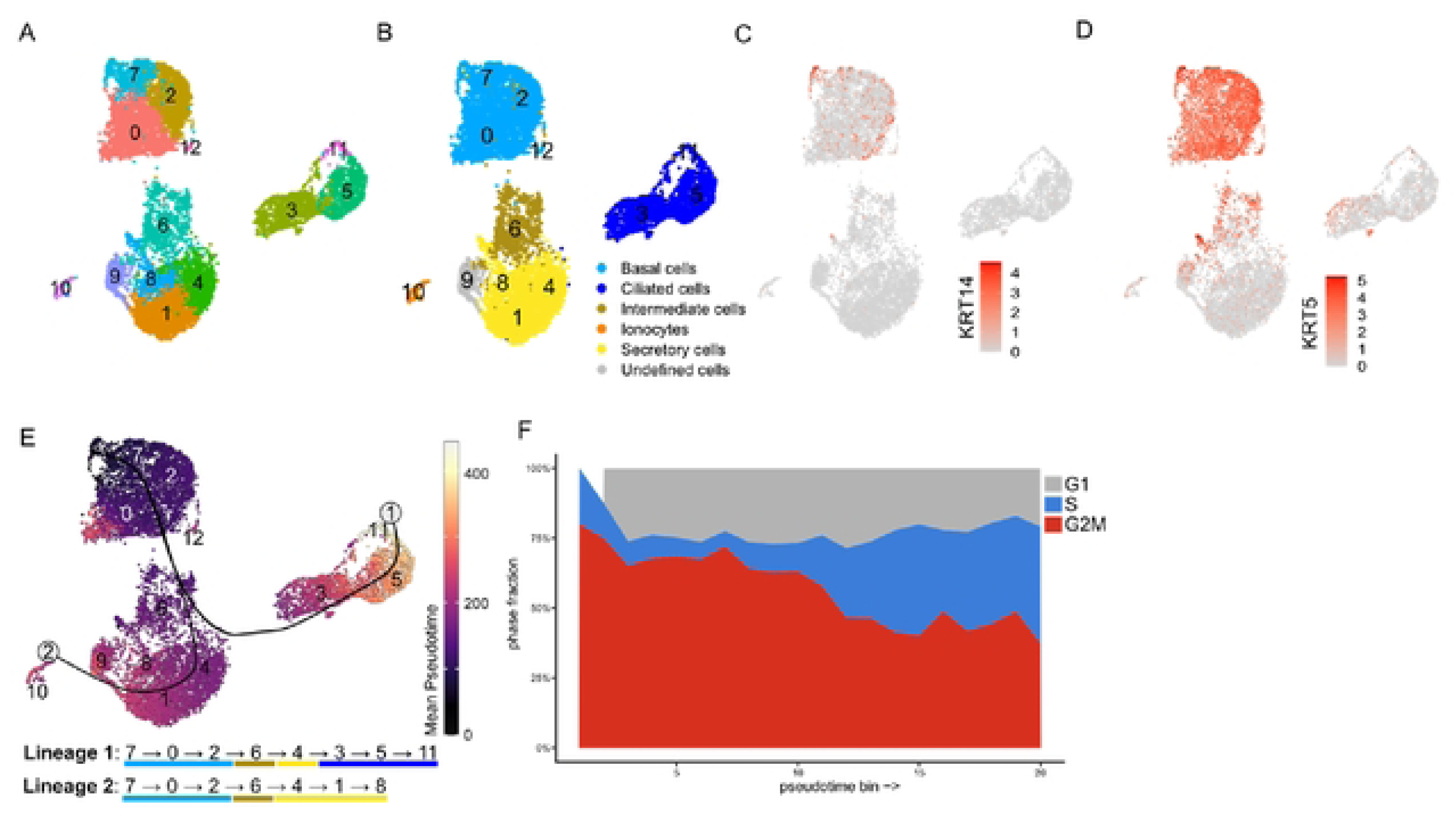
Secretory cells are the precursors of ciliated cells. **A)** Uniform Manifold Approximation and Projection (UMAP) visualization of scRNAseq data from pooled mock-infected BEpC from donors 1-3. The identified clusters are highlighted. **B)** Cell subtype allocation was based on the expression of canonical subtype-specific markers, with a minimum of three markers required for cluster allocation. Basal cells (light blue): TP63, KRT5, DAPL1, KRT15, ITGA6, KRT17; Secretory cells (yellow): SCGB3A1, SCGB1A1, MUC5B, MUC5AC, MUC20, MUC1, SPDEF, BPIFB1, AGR2; Ciliated cells (dark blue): FOXJ1, CAPS, TP73, CCDC78; lonocytes (orange): CFTR, ASCL3, FOXI1, ATP6V1C2. Intermediate cells (brown): clusters expressing markers of basal and secretory cells. Undefined cells (grey): clusters not assignable to known epithelial subtype. **C)** - **D)** UMAP visualization of single-cell transform (SCTransform) -normalized expression of C) KRT14 and D) KRT5. **E)** Trajectory analysis learned with slingshot trajectory interference analysis. Run on the 30-dimensional Harmony embedding and rooted at cluster 7. Harmony UMAP coloured by mean pseudotime, slingshot curves overlaid. Lineages 1 and 2 are reported as the order of clusters they run by. **F)** Cell-cycle phase percentage across pseudotime (lineage mean).

In parallel, we cross-checked the trajectory analysis result using the trajectory inference monocle3 approach, which runs on the Harmony-derived uniform manifold approximation and projection (UMAP) (*39*). To determine the full trajectory analysis, one connected graph was set across the fragmented UMAP islands. As a result, the monocle3 analysis agreed with the ordering of the slingshot analysis and passed through the secretory cell population (cluster 4) to the ciliated cells (Supp. Fig. 2C-D). Collectively, these data highlight that the two approaches agree on the ordering and the trajectory is robust to method choice.

Lastly, to determine a correlation between pseudotime and cell-cycle phases, we mapped the defined phases against pseudotime and found an increase in S phase and a decrease in the G2M phase with pseudotime (Fig. 2F). This correlates with the known promotion of S phase in ciliated cells through MYB factors (*40*, *41*).

Collectively, using the trajectory analysis, we show that the *in vitro* cultured cells differentiate like *in vivo* cells through basal stem cells to secretory cells before maturing to ciliated cells.

### Infected secretory cells are not miscategorized ciliated cells

To assess whether the annotated infected secretory cells could potentially represent misclassified ciliated cells, we examined the expression of canonical secretory and ciliated cell markers within the populations of interest (Fig. 3A). The analysis confirmed that the infected secretory cells do not express canonical mature ciliated cell markers while expressing secretory cell markers, supporting their correct assignment as secretory cells. To determine where the cells would group in a non-infected setting, we next projected the infected cells as a query on the reference mock-treated samples using the FindTransferAnchors and MapQuery approaches (Fig. 3B-E) (*42*). We found that 89.76% of IAV-infected ciliated cells anchored to mock ciliated cells, 94.69% of IAV-infected secretory cells anchored to mock secretory cells, and 95.59% of bystander secretory cells also anchored to mock secretory cells (Fig. 3B). This classification is underpinned by mapping the queries on the mock UMAP graphs (Fig. 3C-E). Additionally, to assess whether a cell subtype misclassification or an infection-induced cell subtype transition could be detected through changes in the relative abundance of the cell subtypes, irrespective of individual cell infection status, we compared the percentages of ciliated and secretory cells between mock and infected samples (Fig. 3F). We did not observe a shift in abundance for either cell subtype.

**Figure 3:**
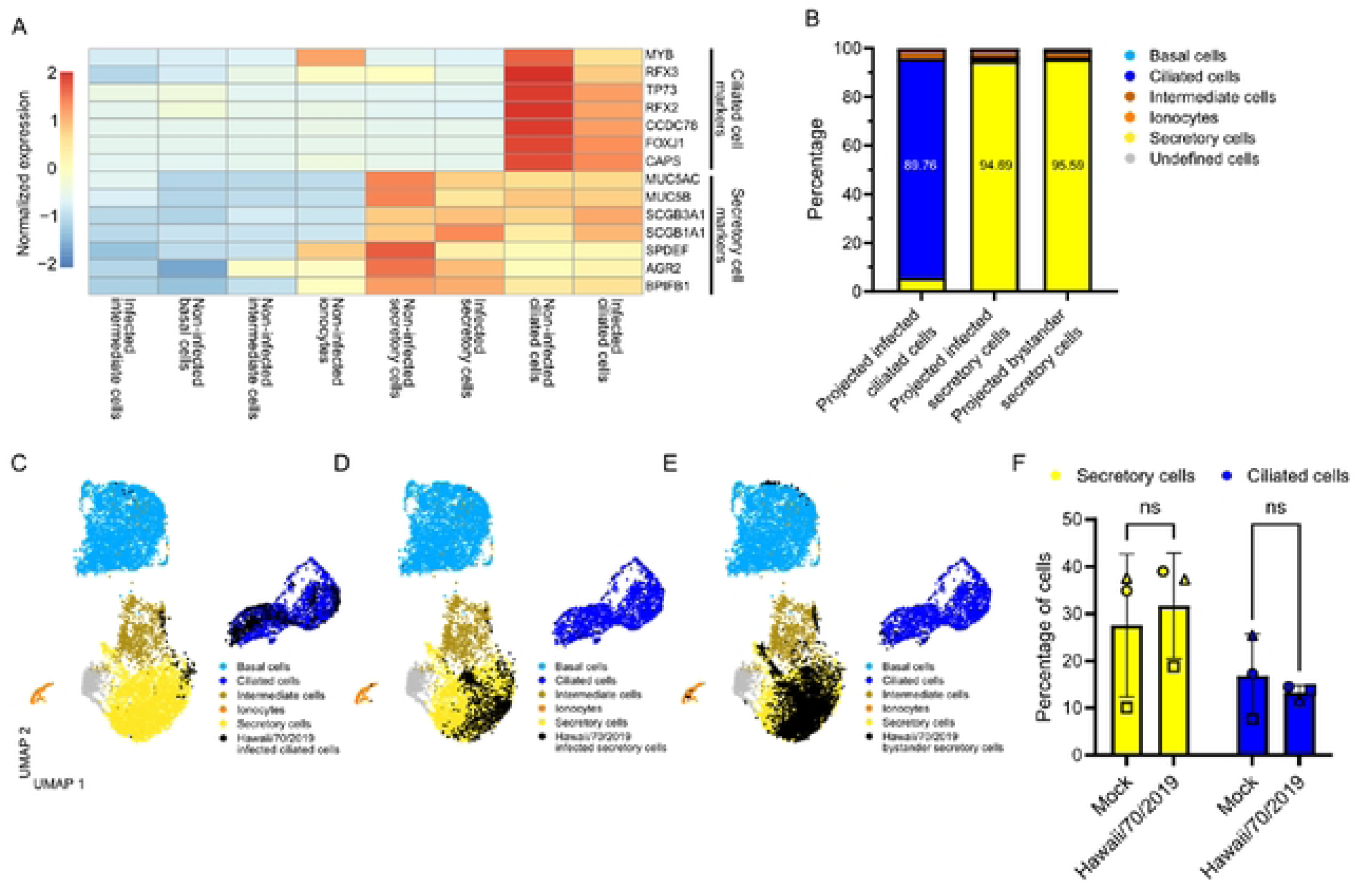
Infected secretory cells are not misclassified ciliated cells. **A)** Pooled BEpC from donors 1-3 infected with H1N1 Hawaii/70/2019 for 6 hours with an MOI of 1. Average expression of ciliated and secretory canonical markers in the Hawaii/70/2019 infected sample (row-scaling). **B)** Projection using Seurat FindTransferAnchors analysis of infected ciliated cells, infected secretory cells, and bystander secretory cells on mock-infected Harmony dataset. **C)** - **E)** Mapped projection of infected ciliated cells, infected secretory cells, and bystander secretory cells using Seurat MapQuery function. Reference mock-infected cells are shown in colours according to cell types, and infected ciliated cells are overlaid in black in C), infected secretory cells are overlaid in black in D), and bystander secretory cells are overlaid in black in E). **F)** Relative cell type composition of BEpC infected with H1N1 Hawaii/70/2019 or mock-infected. Data from donors 1-3 are distinguished by symbol shape: donor 1 (square), donor 2 (triangle), donor 3 (circle). In F), statistical significance was determined using two-way ANOVA with Sidak’s multiple comparisons test. *P≤ 0.05. ns non-significant.

Overall, this further supports our classification of the infected secretory cells as a specific subset of secretory cells, rather than miscategorized ciliated cells, and thus weakens the hypothesis of an infection-induced transition within the initial infection round (Fig. 1I, right panel).

### Cilium-related Gene Ontology terms are overrepresented in infected secretory cells independent of infection-induced changes

To distinguish infection-induced and infection-independent transcript overrepresentation, we defined infection-induced changes as any DEG in the analysis comparing all cells from the Hawaii/70/2019 infected sample over the mock-treated cells (Fig. 4A). The Hawaii/70/2019 infected sample cell population included both infected and bystander cells. Any DEG identified in this comparison was labelled as an ‘infection-induced’ transcriptional alteration and was excluded from the analysis comparing infected over bystander ciliated cells. Following this filtering step, a total of 84 hits remained overrepresented in the infected ciliated cell population, whereas 150 hits were overrepresented in the bystander cell population. Processes related to protein translation were still prominent in the cells primed for infection and independent of any infection-induced changes (Fig. 4B-C). We repeated the same exclusion analysis to compare the hits overrepresented in infected secretory cells in the absence of infection-induced changes (Fig. 4D). This decreased the number of overrepresented hits from 953 to 784 hits. The gene ontology enrichment analysis revealed that the cilium related terms remain overrepresented even after subtracting any infection-induced changes.

**Figure 4:**
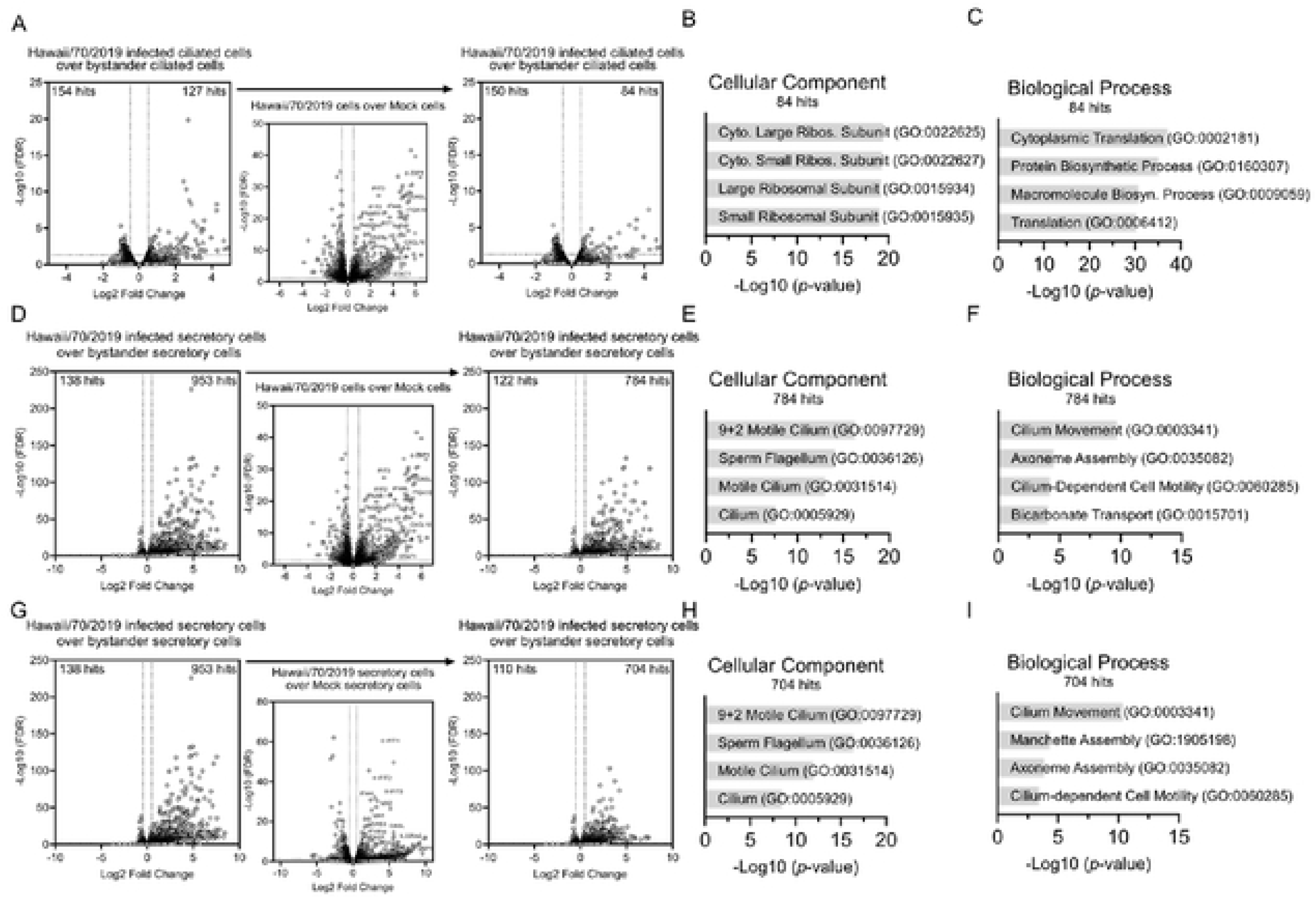
Infected secretory cells are transcriptionally distinct independent of infection-induced changes. **A)** Volcano plot of scRNAseq data from BEpC from donors 1-3 infected with H1N1 Hawaii/70/2019 for 6 hours with an MOI of 1. The scRNAseq dataset was pooled (n=3) to generate pseudobulk expression profiles, and a differential expression analysis was performed of Hawaii/70/2019 infected ciliated cells (686 cells) over bystander (non-infected) ciliated cells (959 cells). Any hit also overrepresented (Log2 (fold change) ≥ ±0.5 and FDR ≤ 0.05) in the Hawaii/70/2019 all cells over mock all cells is excluded from the analysis. **B)** - **C)** Gene ontology (GO) enrichment analysis of the hits overrepresented (Log2 (fold change) ≥ +0.5 and FDR ≤ 0.05) in the Hawaii/70/2019 infected ciliated cells over bystander ciliated cells analysis, filtered for hits also overrepresented (Log2 (fold change) ≥ ±0.5 and FDR ≤ 0.05) in the Hawaii/70/2019 all cells over mock all cells analysis. Top 4 GO terms are shown. **D)** The scRNAseq dataset was pooled (n=3) to generate pseudobulk expression profiles, and a differential expression analysis was performed of Hawaii/70/2019 infected secretory cells (828 cells) over bystander secretory cells (3538 cells). Any hit also overrepresented (Log2 (fold change) ≥ ±0.5 and FDR ≤ 0.05) in the Hawaii/70/2019 all cells over mock all cells is excluded from the analysis. **E)** - **F)** Gene ontology (GO) enrichment analysis of the hits overrepresented (Log2 (fold change) ≥ +0.5 and FDR ≤ 0.05) in the analysis Hawaii/70/2019 infected secretory cells over bystander secretory cells, filtered for hits also overrepresented (Log2 (fold change) ≥ ±0.5 and FDR ≤ 0.05) in the Hawaii/70/2019 all cells over mock all cells analysis. Top 4 GO terms are shown. **G)** The scRNAseq dataset was pooled (n=3) to generate pseudobulk expression profiles, and a differential expression analysis was performed of Hawaii/70/2019 infected secretory cells over bystander secretory cells. Any hit also overrepresented (Log2 (fold change) ≥ ±0.5 and FDR ≤ 0.05) in the Hawaii/70/2019 secretory cells over mock secretory cells is excluded from the analysis. **H)** - **I)** GO analysis of the hits overrepresented (Log2 (fold change) > +0.5 and FDR ≤ 0.05) in the Hawaii/70/2019 infected secretory cells over bystander secretory cells analysis, filtered for hits also overrepresented (Log2 (fold change) ≥ ±0.5 and FDR ≤ 0.05) in the Hawaii/70/2019 secretory cells over mock secretory cells analysis. The top 4 GO terms are shown.

To test whether the overrepresentation of cilium-related transcripts reflects an incomplete exclusion of infection-induced transcriptional differences, and thus not a pre-existing cellular state but rather a cellular response to infection, we applied a more secretory cell-specific definition of infection-induced genes: We excluded all genes identified as differentially expressed in the pseudobulk comparison of the secretory cells from the Hawaii/70/2019 infected samples over the mock secretory cells. Again, the secretory cells from the Hawaii/70/2019 infected samples included both infected and bystander cells. Due to cell subtype-specific infection-induced transcriptional responses (discussed in (*35*)), this approach refined the analysis and further reduced the number of genes overrepresented in infected secretory cells from 784 to 704 (Fig. 4G). Nevertheless, even with this updated filter, the GO analysis identified the same enrichment for cilium-related terms. Therefore, this argues further in favour of a cell-state driven cellular tropism and against an infection-induced cellular transition (Fig. 4H-I).

### IAV infection does not upregulate the canonical ciliogenesis pathway

To assess whether viral infection induces an upregulation of the canonical ciliogenesis program, we analysed the expression of all major genes involved in canonical ciliogenesis (Fig. 5A). Canonical ciliogenesis is initiated by the downregulation of key transcription factors. These are NOTCH1, NOTCH2, and GEMININ (encoded by *GMNN*). We compared secretory cells from the infected samples (independent of the infection status of the cells) to mock secretory cells. However, none of these three regulators are differentially expressed in secretory cells from infected versus mock-treated samples, arguing against an activation of the canonical ciliogenesis pathway following infection (Fig. 5B right panel, bold). Next, we assessed the expression of transcripts that are induced upon inhibition of these transcriptional regulators to promote ciliogenesis (Fig. 5A). Among these, only *GMNC*, which encodes GEMC1 proteins, was significantly upregulated in secretory cells of the infected samples (Fig. 5B left panel), whereas all other ciliogenesis genes showed no differential expression (Fig. 5B right panel), arguing further against an activation of the canonical ciliogenesis pathway following infection.

**Figure 5:**
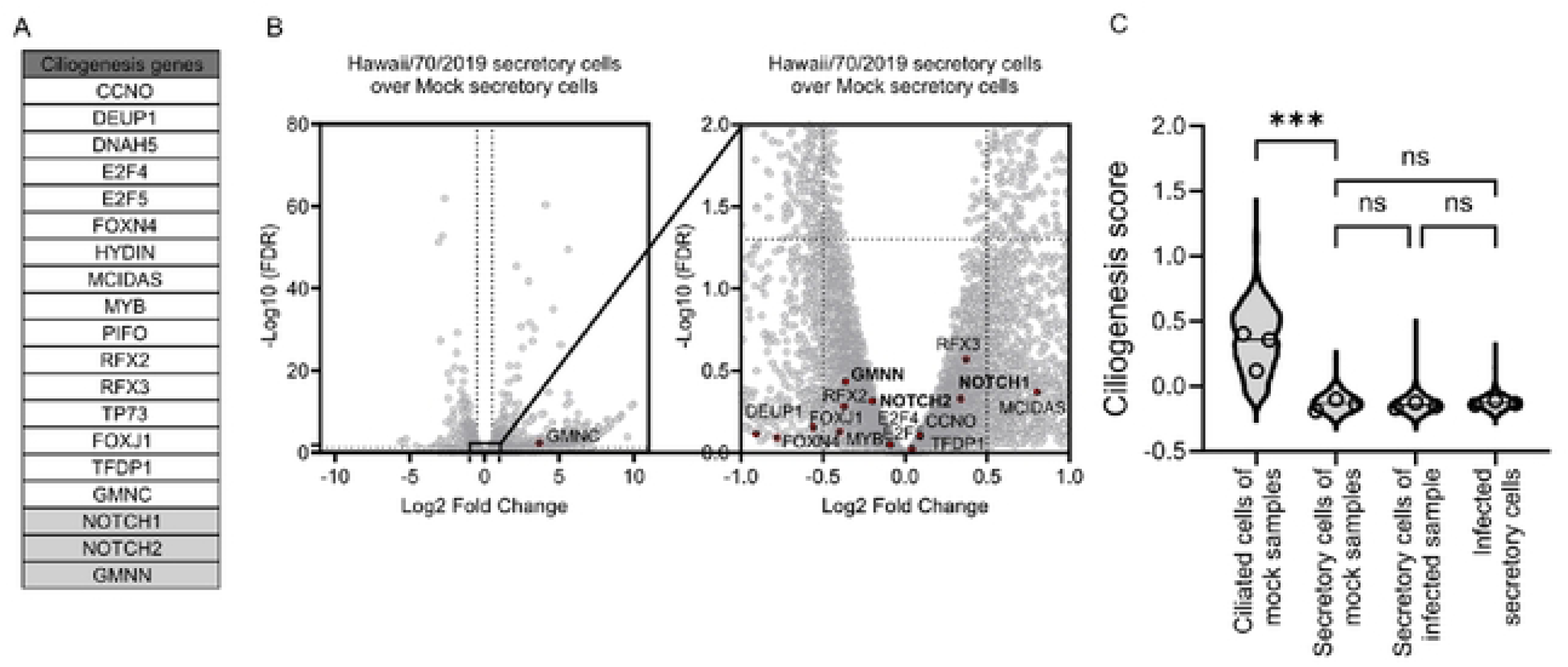
Influenza A virus infection does not induce the canonical ciliogenesis pathway at 6 hours post infection. **A)** List of canonical regulators of ciliogenesis. **B)** The scRNAseq data were pooled to generate pseudobulk expression profiles, and a differential expression analysis was performed of Hawaii/70/2019 secretory cells over mock secretory cells. The canonical ciliogenesis genes listed in A) are highlighted. **C)** Canonical ciliogenesis expression scores for each cell were calculated using Seurat AddModuleScore analysis. Only the genes upregulated during ciliogenesis were included (exclusion of GMNN, NOTCH1, and NOTCH2). Shown are the ciliogenesis scores from mock ciliated cells, mock secretory cells, secretory cells from the infected samples, and infected secretory cells. In C), statistical significance was calculated on the average expression of each BEpC donor, using two-way ANOVA with Tukey’s multiple comparisons test. *P≤0.05, **P≤0.01, ***P≤ 0.001. ns non-significant.

To further determine at a single-cell level whether secretory cells display an increase in canonical ciliogenesis upon infection, we moved beyond the pseudobulk approach, which averages gene expression across all cells. We instead investigated expressional changes at a single-cell level by defining a ciliogenesis signature score for each individual cell (*43*). This signature score encompassed the main genes upregulated during canonical ciliogenesis and excluded the downregulated factors (Fig. 5A) (i.e., *GMNN, NOTCH1,* and *NOTCH2*). The mock ciliated cell population was included as a positive control of the signature score (Fig. 5C). Using this single-cell-resolved approach, we revealed that there is no significant increase in the overall canonical ciliogenesis score between mock secretory and secretory cells from the infected sample, independent of the infection status of the cells, nor is there an increase in the ciliogenesis score between mock secretory cells versus infected secretory cells. Moreover, none of these factors strongly correlate positively with viral HA segment expression in infected secretory cells (Supp. Fig. 3-4), instead, most markers show no or slightly negatively correlation with increased viral replication. Collectively, these results highlight that the infection of secretory cells as such does not increase canonical ciliogenesis within 6 hours post-infection.

In summary, our findings support the hypothesis that IAV tropism is driven by a pre-existing secretory cell state (Fig. 1I, left panel), rather than by an infection-induced cell state switch (Fig. 1I right panel).

## DISCUSSION

Overall, our study demonstrates that within the first round of infection of the human airway epithelium, seasonal H1N1 IAV infects a subcluster of secretory cells with a distinct transcriptome compared to the transcriptomes of bystander secretory cells. The infected secretory cells show significant overexpression of cilium-related genes. We further show that the *in vitro* cell cultures differentiate as *in vivo* cells, whereby the basal stem cells differentiate through secretory cells to mature ciliated cells. We tested and confirmed that the infected secretory cells are not misclassified ciliated cells despite the overrepresentation of cilium-related genes. Furthermore, using an *ex post facto* study approach, we show that the infection itself does not increase the canonical ciliogenesis pathway and therefore affirm the hypothesis that seasonal H1N1 IAV preferentially infects secretory cells that are primed for ciliogenesis.

A major limitation of this study is the *ex post facto* design, which limits our ability to conclusively define the complete set of infection-induced changes. Consequently, we cannot fully exclude the possibility that the observed cell states reflect, at least in part, an infection-induced transition rather than a pre-existing cellular state. Previous studies, including Reffsin *et al.* (2026), tried to overcome these limitations posed by *ex post facto* approaches by implementing a retrospective single-cell method (*44*). Such approaches have great potential for profiling susceptible cells and remodelling the cell state prior to infection. Nonetheless, the approach presented by Reffsin *et al.* (2026) is dependent on genetic manipulation of cells, an experimental approach that is still not broadly applicable to primary air-liquid interface cultures such as those used in our study. Additionally, the approach is dependent on cell division, which is considerably more challenging in differentiated cells compared to immortalized cell line cultures, such as those used by Reffsin *et al.* (2026). Hence, while retrospective single-cell methods hold great promise for future studies in our field, their implementation in differentiated primary epithelial cells remains limited.

Our study was conducted at a single time point of 6 hours post-infection. Consequently, we analysed an early cell state switch but did not seek to identify a complete transition from secretory to ciliated cells. Based on previous literature, this process is known to take several days (*45*, *46*). Thus, we aimed to determine if there was an initial activation of the ciliogenesis pathway. To strengthen this analysis, it would be interesting to follow up at a later point of infection to assess changes in the cell type composition. Such analysis could potentially better resolve whether the virus promotes an infection-induced transition to ciliated cells at later time points post infection. Potential anchor points could be the results of Matrosovich *et al.* (2004) and Kelly *et al.* (2022), which both showed that H1N1 influenza viruses preferentially infect non-ciliated and secretory cells, respectively, during the initial infection cycle, but upon multicycle replication, the viral staining was also observed in ciliated cells (*47*, *48*). If these observations were to be combined with a quantitative analysis of the cell populations in question, one could potentially determine if the infected ciliated cells arise via an infection-induced transition from secretory cells. Support for the ability of influenza viruses to induce cellular differentiation is provided by the study by Hou *et al.* (2012) (*49*). They found that a variety of IAVs induce, within 18 hours of infection, a monocyte differentiation into mature dendritic cells.

While our study focused on identifying a potential cell subpopulation tropism, it did not investigate the underlying processes driving such a preferential targeting. As discussed above, Greaney *et al.* (2023) identified SARS-CoV-2 infecting progenitors of ciliated cells. The authors hypothesized that the virus achieves greater overall infection output when taking advantage of the rapid epithelial turnover rate in progenitor cells compared to mature ciliated cells (*32*). Regarding IAV infections, Roach *et al.* (2024) identified secretory cells showing higher viral replication levels during the initial IAV infection round, a finding that was also supported by our dataset (*35*). On the other hand, Roach *et al.* (2024) identified ciliated cells being associated with increased viral production during multi-cycle replication, while secretory cells were linked with an overall reduced virus output (*50*). Together, these studies, in combination with our observation of a tropism for ciliogenesis-primed secretory cells during the initial infection round, support a model whereby IAVs target specific cell subpopulations to maximize their viral output during subsequent replication rounds. This finding would also positively correlate with the stronger antiviral immune response in secretory cells compared to ciliated cells, identified by Roach *et al.* (2024) and confirmed in our previous analysis of the same dataset as presented above (*35*). Collectively, these findings suggest that the preferential targeting of ciliogenesis-primed secretory cells may represent a strategy that allows IAV to optimize viral replication output.

Overall, this study sought to determine whether IAV infection is driven by a pre-existing secretory cell state (i.e. cell-state-driven tropism) or if infection itself induces a transition in secretory cell state (i.e. infection-induced cell state transition). Due to the limitations of *ex post facto* approaches, we cannot fully exclude the latter. However, our findings strongly support the former hypothesis and thus, refine IAV tropism beyond cell type to encompass epithelial differentiation state.

## METHODS

### Single-cell RNA sequencing processing of infected primary human bronchial cultures

The single-cell RNA sequencing dataset generation and analysis were previously described in von Kempis *et al.,* 2026 (*35*).

### Single-cell RNA sequencing SUSHI pipeline

The single-cell RNA sequencing data analysis used the SUSHI framework (*51*) and was previously described in von Kempis *et al.* (2026) (*35*). In brief, it includes the following analysis. A chimeric reference genome was constructed using the R package ezRun (v3.19.1) (*52*) by including the human genome assembly (GRCh38.p13, GENCODE Release 42) (*53*) with the nucleotide sequences of the IAV strain H1N1 A/Hawaii/70/2019. Using 10x Genomics Cell Ranger v7.2.0, we performed sample demultiplexing, alignment of reads against this chimeric reference, and generation of feature–barcode count matrices (*54*). We enabled pre-mRNA counting (include-introns=true) to also capture unspliced transcripts. Downstream analysis was performed on the resulting count matrices (R package Seurat v5.1.0 under R version 4.4.0 and Bioconductor 3.19) (*55*). DecontX (celda v1.20.0) and SoupX (v1.6.2) were used to estimate and remove ambient RNA contamination (*56*, *57*). scDblFinder was used to identify and remove doublets (*58*). Low-quality cells were filtered using a median-absolute-deviation approach (*59*). Cells with ribosomal protein gene content exceeding 70% were removed, and genes expressed in fewer than 0.01% of cells were filtered out before downstream analysis. SCTransform v2 (sctransform v0.4.1) was used for data normalization and variance stabilisation (*60*). Principal component analysis (PCA) was performed for dimensional reduction. The Harmony algorithm was used for donor-level batch correction and integration. This was conducted separately for each experimental condition (mock, Hawaii/70/2019-infected) (*61*). Cell clusters were identified using the Louvain algorithm (resolution of 0.6). Cluster marker genes were determined by a Wilcoxon rank-sum test, retaining genes with an average log₂(fold-change) ≥ 0.4, a Bonferroni-adjusted p-value < 0.01, expressed in at least 10% of cells in the tested cluster, and with a minimum difference in detection rate of 10% between the tested cluster and all other cells. Epithelial cell subtype allocation was based on the expression of canonical epithelial cell subtype-specific markers, with a minimum of three canonical markers required for cluster allocation. Basal cells: TP63, KRT5, DAPL1, KRT15, ITGA6, KRT17; Secretory cells: SCGB3A1, SCGB1A1, MUC5B, MUC5AC, MUC20, MUC1, SPDEF, BPIFB1, AGR2; Ciliated cells: FOXJ1, CAPS, TP73, CCDC78; Ionocytes: CFTR, ASCL3, FOXI1, ATP6V1C2. Intermediate cells: clusters expressing markers of basal and secretory cells. Undefined cells: clusters not assignable to known epithelial subtype. Infected cells were defined as those expressing at least four viral gene segments, with a segment considered expressed if its unique molecular identifier (UMI) count exceeded 1.5 times the mean expression of that segment across all cells in the same sample. Per-cell viral transcript abundance was computed as log10(1 + the sum of viral UMIs) and summarised per cell type as the mean of these per-cell values within each group. The data are visualised using GraphPad Prism version 11.0 and RStudio Server 2026.05.0.

### Pseudobulk dataset

Pseudobulk differential expression analysis was performed using the SUSHI framework (*51*) and was previously described in von Kempis *et al.* (2026) (*35*). In brief, each sample was treated as a biological replicate. Pseudobulk count matrices were generated by summing raw UMI counts per cell type per sample from the individually processed Seurat objects. Differential expression between conditions was tested using DESeq2 (*62*) under R version 4.4.0 and Bioconductor 3.19, with the design formula ∼ donor + group. A differentially expressed gene (DEG) was defined with an absolute log2 fold change ≥ 0.5 and a false discovery rate (FDR) ≤ 0.05. Gene ontology (GO) enrichment of DEGs was performed using Enrichr (*63–65*).

### Trajectory analysis

Trajectories were inferred *via* the R Bioconductor package ‘slingshot’ (*36*) and the ‘Monocle3’ package (*39*). Pseudotime was estimated primarily with slingshot, run on the 30-dimensional Harmony-corrected embedding with Seurat cluster labels as lineage nodes. The mock trajectory was rooted at the basal cluster (highest KRT5/KRT14); the infected trajectory was restricted to the six clusters spanning the differentiation axis (resolution 0.4, excluding basal and ionocyte clusters) and rooted at the early-secretory cluster, yielding three lineages: An intermediate spur, the ciliated backbone, and an infected-secretory spur. Principal curves were fitted in Harmony space and projected onto the UMAP for visualisation only. Per-cell pseudotimes and curve weights were taken from slingPseudotime and slingCurveWeights. As a method-independent check, the same trajectory was learned with monocle3 on the identical cell set and embedding, via the ‘cluster_cells’, ‘learn_graph’ (use_partition = FALSE), and ‘order_cells’ functions, rooting at the principal-graph vertex nearest the majority of cells in the same root cluster used for slingshot. Agreement between the two orderings was assessed by per-cell correlation of slingshot and monocle3 pseudotime. Genes varying along the principal graph were identified with graph_test(neighbor_graph = "principal_graph") at Moran’s I > 0.1 and q < 0.05. Analyses were performed in R 4.6 with Seurat v5.

### Scoring analysis

To assign the ciliogenesis scores to the single-cell RNA sequencing dataset described above, we used the Seurat AddModuleScore function on RNA data with default settings and the genes listed above (*43*). Analyses were performed in R 4.6 with Seurat v5.

### Projection analysis

To project the cells from the infected samples onto the Harmony object of the mock-treated cells, we used the Seurat FindTransferAnchors and MapQuery functions (*66*). The projection was based on the 3000 most variable features of the mock object (VariableFeatures assay). The projection used the first 30 principal components as a reference reduction and a log normalization. Analyses were performed in R 4.6 with Seurat v5.

## DATA AVAILABILITY

All data needed to evaluate the conclusions in the paper, except for the single-cell sequencing data, are present in the paper and/or the Supplementary Materials. Single-cell sequencing data have been deposited in the NCBI Gene Expression Omnibus repository under accession number GSE328834.

## ACKNOWLEDGEMENTS

We thank the Functional Genomics Center Zurich for their support with the single-cell transcriptomics approach, sequencing, and data analysis. This study was funded by the Swiss National Science Foundation (grant number 310030_204166 to SSt) and the UZH Candoc Grant (grant number FK-25-039 to JvK).

